# Sex differences in gene regulation and its impact on cancer incidence

**DOI:** 10.1101/2025.11.21.689715

**Authors:** Camila M. Lopes-Ramos, Rebekka Burkholz, Marouen Ben Guebila, Viola Fanfani, Enakshi Saha, Katherine H. Shutta, Kimberly Glass, John Quackenbush, Dawn L. DeMeo

## Abstract

Sex differences in the incidence rates of many cancer types are well-documented. While differences in lifestyle, environmental exposures, and hormone levels can influence disease risk and incidence, there is limited understanding of how cancer driver genes are differentially regulated between males and females in normal tissues, and how these differences may contribute to the observed sex differences in cancer epidemiology. We studied 8,279 gene regulatory networks across 29 non-cancerous tissues in the GTEx dataset, referred here as normal tissues. Using network centrality measures, we compared the networks between males and females, focusing on interactions between 644 transcription factors and 476 cancer genes (COSMIC Cancer Gene Census). Cancer genes were differentially targeted by transcription factors in males and females across normal tissues. We found an overrepresentation of sex-biased cancer genes on the X chromosome, particularly among genes escaping X chromosome inactivation, with higher regulatory targeting of tumor suppressor genes in females compared to males. Key signaling pathways, including WNT, NOTCH, and p53, showed differential transcriptional targeting by sex. We observed higher targeting of cancer-related pathways in females for tissues that have higher tumor incidence in females (breast, lung, and thyroid) and higher targeting in males for tissues with increased tumor incidence in males (stomach, colon, and liver). Sex-biased transcription factors that were consistently observed across multiple tissues are enriched for sex hormone response elements in their promoters. Our findings showed that normal tissues have sex-biased regulation of genes implicated in tumorigenesis, helping to explain the molecular basis of sex differences observed in cancer incidence rates and in identifying targets for cancer prevention in a sex-aware manner.

## Introduction

There are significant differences in cancer risk and incidence rates between males and females^1–4^. Males tend to have higher incidence rates for most cancer types, including those of the gastrointestinal tract, kidney, and bladder. However, females have higher incidence rates for breast, thyroid, and lung adenocarcinoma cancers. A combination of factors may contribute to differential disease risk and incidence between males and females, including lifestyle and environmental exposures, metabolism, steroid hormones, and sex chromosome complement^5^. Genomic studies have shown that tumors exhibit distinct molecular profiles between males and females that influence disease progression, treatment response, and clinical outcomes in a sex-dependent manner^6–8^. Despite these advancements, a knowledge gap persists in understanding how cancer driver genes are differentially regulated between males and females in normal tissues, and how these differences may contribute to the observed sex differences in cancer epidemiology.

Gene regulatory networks capture the complex interplay between transcription factors and their target gene expression. We have shown that constructing gene regulatory networks for each sex can identify sex-differential transcriptional regulatory processes in health and disease. For instance, in colon cancer, higher transcriptional targeting of drug metabolism genes was associated with improved survival in females receiving chemotherapy but not in males^9^. We also found sex-differential targeting of several oncogenes and tumor suppressor genes in lung adenocarcinoma, including genes relevant to drug treatment, and overall survival^10^. In analyzing 8,279 gene regulatory networks across 29 non-cancerous human tissues, we found that sex-differential transcriptional targeting of genes can occur even when mean gene expression levels are similar between males and females^11^. Such differences in regulatory networks could become important contributors to sex-divergent disease development over time, as the effects of aging and environmental exposures may influence gene networks in sex-specific ways.

In the present study, we analyzed 8,279 gene regulatory networks to investigate the transcriptional regulation of cancer driver genes and cancer-related pathways across 29 non-cancerous, non-reproductive tissues using the GTEx dataset. We hypothesized that comparing networks of normal tissues between sexes may reveal sex-biased transcriptional regulation of established cancer driver genes and pathways. We found higher targeting of pathways with known relevance in cancer in females for tissues with higher tumor incidence in females such as breast, lung, and thyroid. Conversely, we observed higher targeting of cancer-related pathways in males for tissues with higher tumor incidence in males, including stomach, colon, and liver. Sex-biased transcription factors found in multiple tissues were significantly enriched for sex hormone binding motifs within their promoter regions. Through this network-based approach, we provide an effective framework to identify sex-biased regulation of cancer-related genes across normal tissues, thereby enhancing our understanding of the molecular mechanisms underlying sex differences in cancer incidence.

## Results

### Tumor suppressor genes are highly targeted in females

We studied gene regulatory networks across 29 non-cancerous tissues from the GTEx dataset, hereafter referred to as normal tissues (**Figure 1; Table S1)**. These networks provide a regulatory map between 644 transcription factors and 30,243 target genes, with the strength of regulation represented by edge weights. For each of the 29 tissues, we identified genes that demonstrated sex-biased differential targeting by comparing the edge weights between male and female subjects, as reported in^11^.

**Figure 1.**
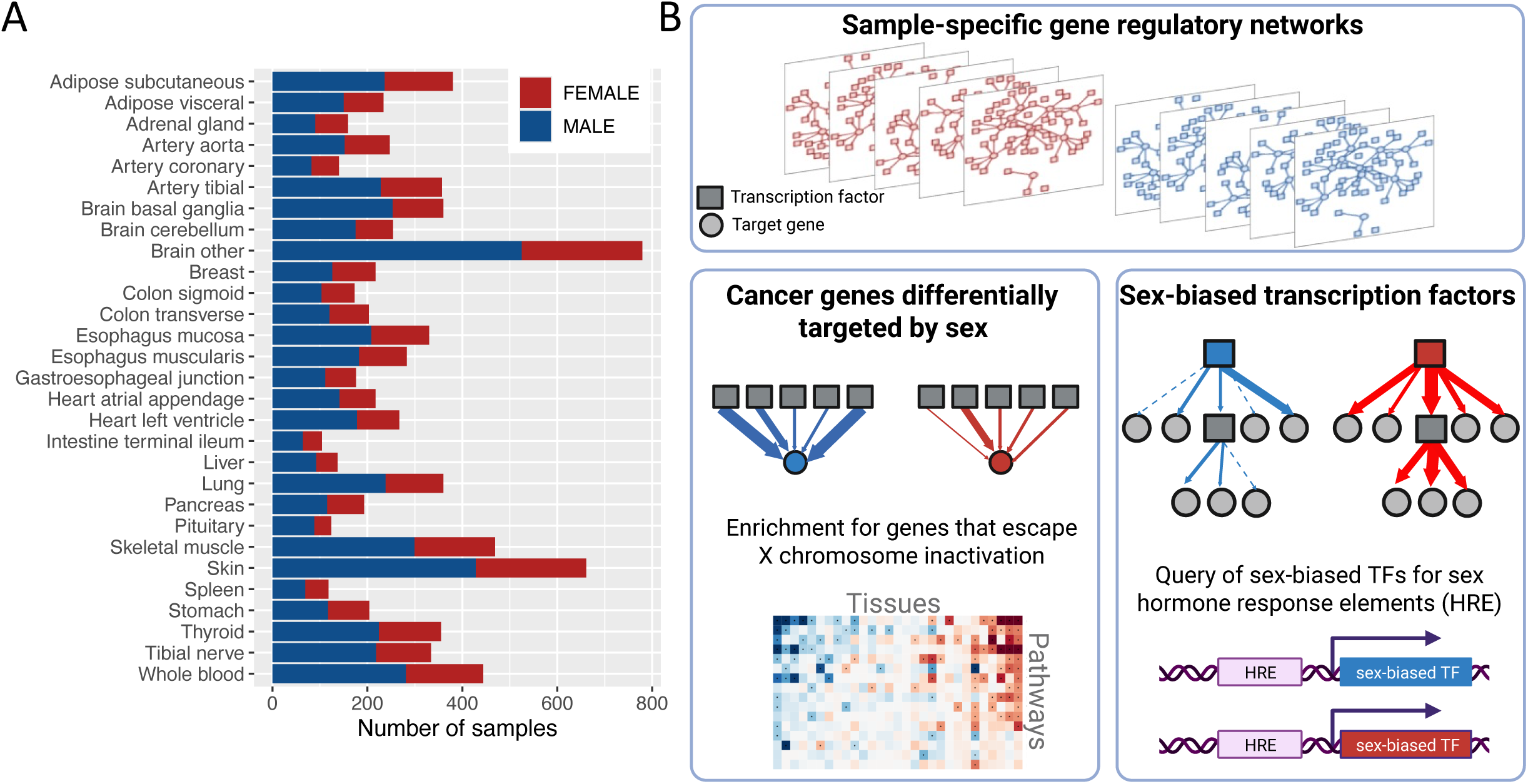
Study design overview. A) Number of female (red) and male (blue) samples analyzed from the GTEx dataset for each tissue included in the study. B) Schematic summary of the main analytical approach: (top) Sample-specific gene regulatory networks for each individual, capturing transcription factor-gene regulatory relationships; (bottom left) identification of cancer genes differentially targeted by sex and pathway enrichment analyses, including assessment of enrichment for genes that escape X chromosome inactivation; (bottom right) identification of sex-biased transcription factors through cascade modeling, along with enrichment analysis for sex hormone response elements in their promoter regions.

Here we focused on 476 cancer driver genes as defined by the COSMIC Cancer Gene Census v92; 23 of those are on the X chromosome (**Table S2**). In this analytic cancer gene set, cancer genes are not uniformly distributed across the genome. On average, 1.6% of genes across all autosomes are categorized as cancer genes, compared to 2.3% of genes on the X chromosome. Notably, 6% of genes that escape X chromosome inactivation are categorized as cancer genes. We observed that cancer genes tend to have higher in-degree (sum of regulatory edge weights connecting a given gene with transcription factors) and closeness (inverse of the sum of the shortest path distances from a given node to all other nodes), compared to non-cancer genes, indicating that cancer genes might have a stronger influence in regulatory networks compared to non-cancer genes **(Figure S1)**. In addition, we observed an overrepresentation of these cancer genes among the genes that demonstrate sex-biased differential targeting (**Figure 2A**). These cancer genes with sex-biased targeting were not randomly scattered across the genome. We observed an overrepresentation of cancer genes with sex-biased targeting in regions of the X chromosome that escape X chromosome inactivation, compared to other X chromosome regions and autosomes. For this we used the list of escape genes previously reported in ^12^. On average, across all tissues, 80% of cancer genes that escape X chromosome inactivation exhibited sex-biased targeting, while 10% of cancer genes on autosomes had sex-biased targeting (**Figure 2B**).

**Figure 2.**
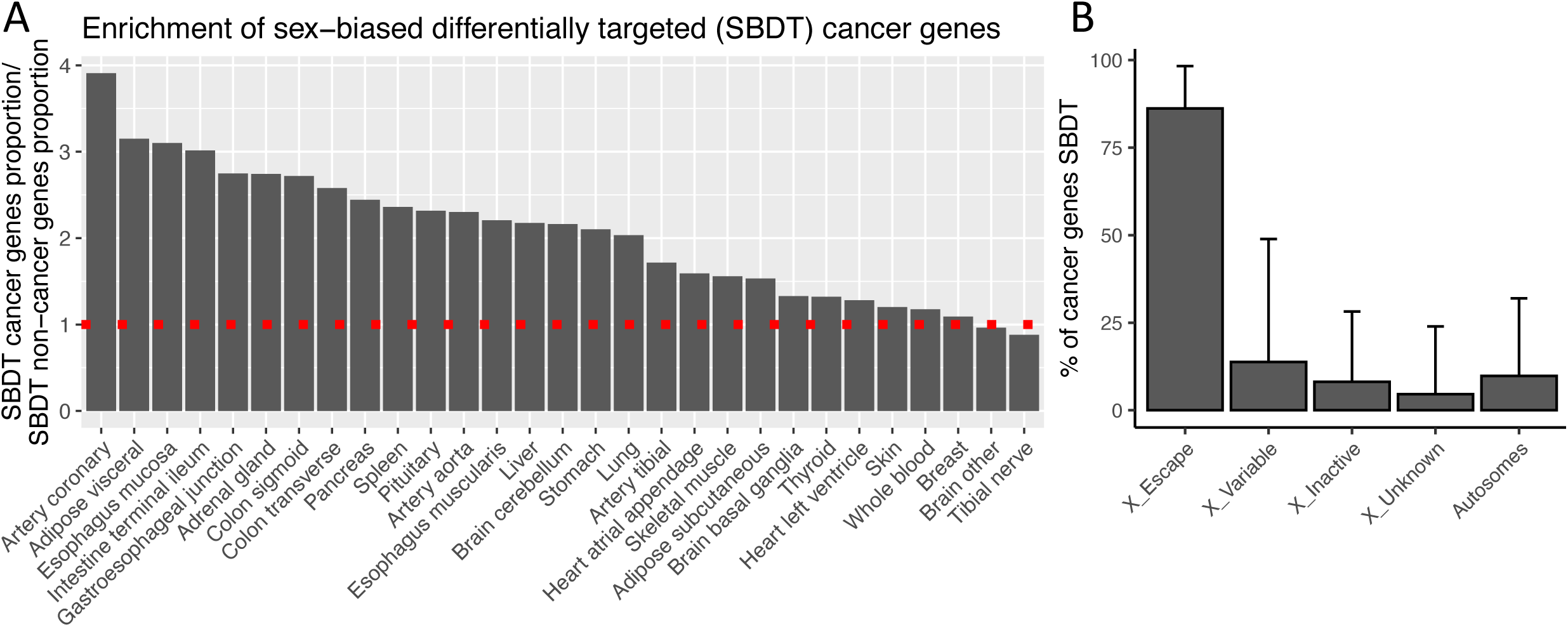
Cancer genes are enriched for sex-biased regulatory edges. A) The bar plot shows the proportion of sex-biased differentially targeted (SBDT) cancer genes to the proportion of SBDT non-cancer genes (SBDT cancer genes/476 total cancer genes)/(SBDT non-cancer genes/29,767 total non-cancer genes). The red line represents where the proportion of SBDT cancer genes equals that of SBDT non-cancer genes. B) The percentage of cancer genes that are SBDT is shown by allosomal and autosomal region, with bars indicating the mean and standard deviation across 29 tissues. The X chromosome is categorized based on X chromosome inactivation status: escape from inactivation, variable escape, inactive, and unknown.

We further evaluated whether cancer genes with sex-biased targeting tended to be associated with higher targeting in females or males. We separately analyzed the cancer genes based on their classification as oncogenes or tumor suppressor genes, as provided by COSMIC. We did not observe a consistent pattern of transcription factor targeting for oncogenes across tissues. However, we found a significant overrepresentation of tumor suppressor genes in the female-biased direction (**Figure 3A-B**). Across all tissues, the median proportion of sex-biased tumor suppressor genes in the female direction was 0.50, compared to 0.17 in the male direction (Wilcoxon rank test, p = 0.007). Three tumor suppressor genes, namely *DDX3X, KDM5C,* and *ZRSR2*, demonstrated differential targeting by sex across all 29 tissues, and *SMC1A* across 27 tissues (**Figure 3C**). These four genes are located on the X chromosome and escape X chromosome inactivation in females.

**Figure 3.**
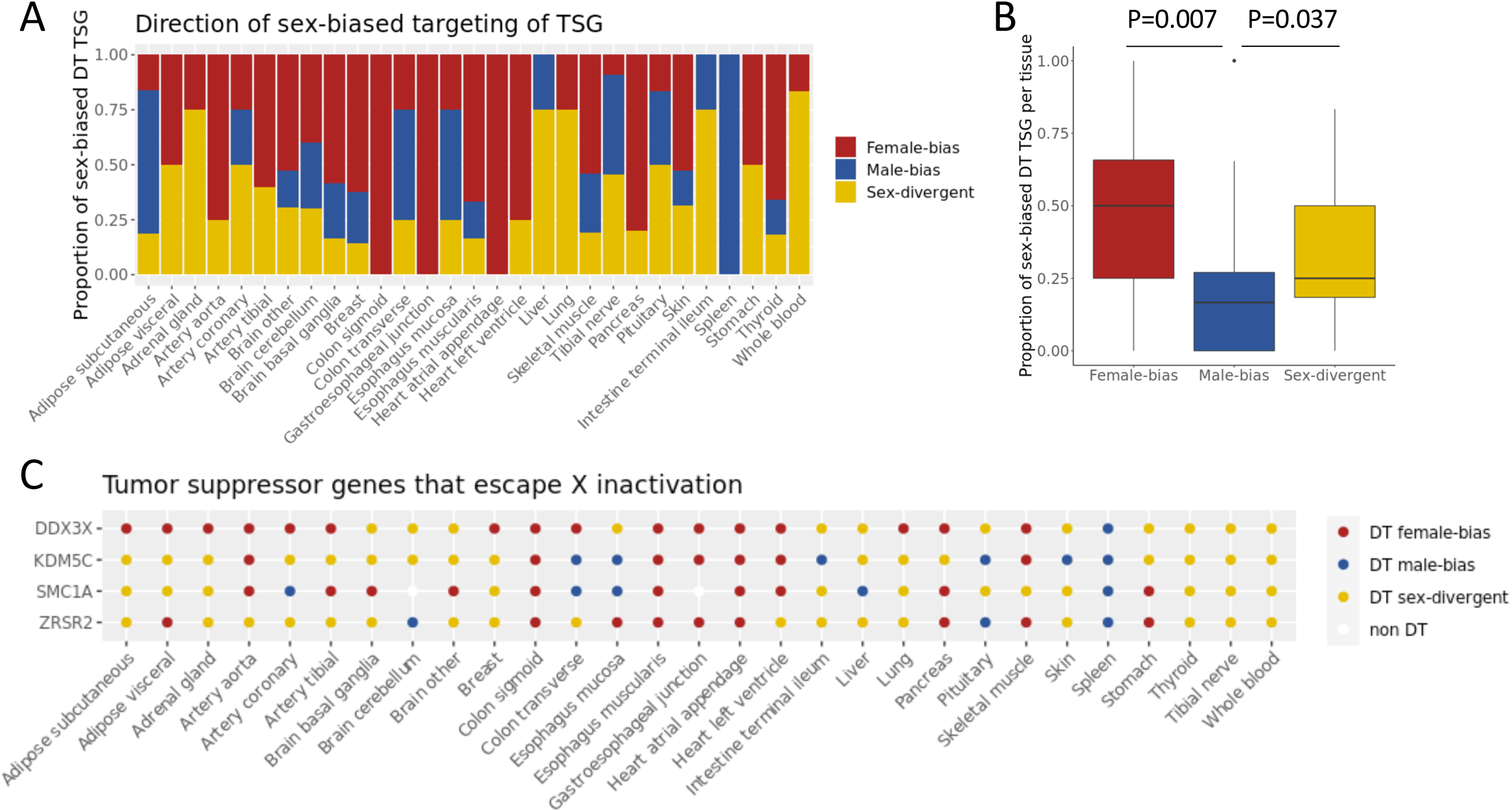
Tumor suppressor genes (TSG) have higher regulatory targeting in females. A) Proportion of TSG that are differentially targeted (DT) by sex. Female-bias: Transcription factor-to-gene regulatory edges are significantly higher in females; Male-bias: transcription factor-to-gene regulatory edges are significantly higher in males; Sex-divergent: genes targeted in both sexes, but by different set of transcription factor. B) Boxplot summarizing the 29 tissues in panel A. P-value calculated by Wilcoxon rank test, two-sided, paired comparing male-bias versus female-bias and male-bias versus sex-divergent. Female-bias versus sex-divergent comparison was not statistically significant. C) TSG that escape X-chromosome inactivation in females are differentially targeted by sex across tissues.

### Cancer pathways are differentially targeted by sex

We performed pre-ranked gene set enrichment analysis across all genes ranked on differential targeting (based on in-degree) by sex. We found that cancer-related pathways are enriched for genes differentially targeted between males and females. The results for all KEGG pathways as well as the leading genes are presented in **Table S3**, while **Figure 4A** provides a heatmap emphasizing the cancer-related pathways. Our analysis revealed a higher targeting of cancer-related pathways in females compared to males in tissues that exhibit a higher incidence of tumors in females, such as breast, lung, and thyroid tissues. Conversely, we observed a higher targeting of cancer-related pathways in males for tissues that have a higher tumor incidence in males, including stomach, transverse colon, and liver.

**Figure 4.**
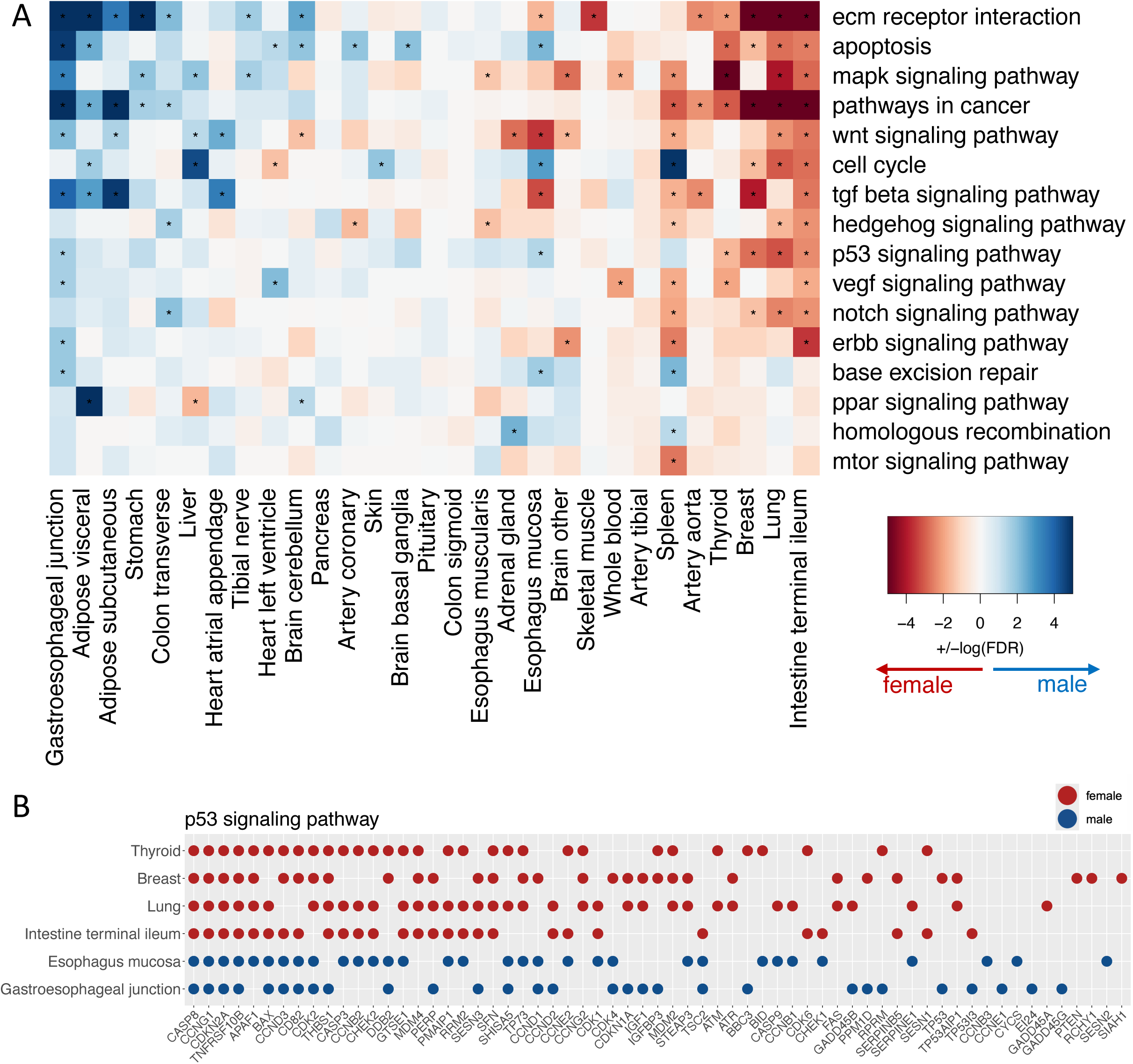
Cancer-related pathways are enriched for genes differentially targeted by sex. A) Gene set enrichment analysis was performed for each tissue using genes ranked by sex-differential targeting. The heatmap shows log_10_(FDR) significance values, with positive values (blue) indicating higher targeting in males and negative values (red) indicating higher targeting in females. Tissues (columns) are ordered by their average significance across all pathways, while pathways (rows) are ordered by the number of tissues in which they achieved FDR<0.05. B) Leading genes identified in the p53 signaling pathways for tissues with FDR<0.05 are shown. Enrichment scores across all pathways and leading genes can be found in Table S2.

The pathways that exhibited sex-biased enrichment across most tissues were the Extracellular Matrix (ECM) Receptor Interaction, apoptosis, MAPK signaling and Pathways in Cancer (designated in the KEGG database as a superset of 318 genes, and part of many other cancer-related and more specific pathways). We identified many examples of sex-biased cancer-related pathways in specific tissues. For instance, the Notch signaling pathway showed female-biased targeting in breast, lung, intestine terminal ileum and spleen, and male-biased targeting in transverse colon. Similar patterns were also observed for WNT, and MAPK signaling pathways. For the p53 signaling pathway, we observed female-biased targeting in thyroid, breast, lung, and intestine terminal ileum tissues. In contrast, we observed male-biased targeting of p53 in esophagus mucosa and gastroesophageal junction. **Figure 4B** highlights the leading genes involved in the p53 pathway for each tissue. Some genes, such as *CASP8*, *CCNG1*, and *CDKN2A*, are consistently identified across all six tissues, while others show tissue-specific targeting patterns. We also observed differences between subregions of the same tissue. For example, in the transverse colon, there was a higher targeting in males for ECM receptor interaction, Hedgehog, Notch signaling, and Pathways in Cancer. However, we did not observe significant sex bias in the sigmoid colon.

Taken together, these analyses revealed sex-biased regulation of cancer pathways across 29 normal tissues. The patterns of sex bias were consistent with known differences in cancer incidence between males and females for different tissue types. **Table S3** provides a comprehensive resource for identifying sex-biased pathways and their leading genes across normal tissues, which can help guide sex-aware searches for cancer prevention targets.

### Sex-biased transcription factors

We used a cascade model to identify transcription factors that drive sex-biased gene regulation (see Methods). The transcription factors were scored based on their likelihood of reaching and influencing genes in the gene regulatory network. Then, we identified the transcription factors that had significantly different scores between males and females. The transcription factor score is positively associated with its out-degree in the network binary prior, which represents the number of genes that have the transcription factor motif and can potentially be regulated by it **(Figure S2)**. In our analyses, transcription factors with higher scores have more starting connections and a higher chance to regulate more genes. In contrast, we observed that sex-biased transcription factors have lower out-degrees in the prior, indicating that they target fewer genes and are not hubs in the network **(Figure S2)**.

On average, across 29 tissues, we identified 10 female-biased and 10 male-biased transcription factors **(Figure 5A)**. The liver and pancreas had the most, with 49 sex-biased transcription factors each. These analyses suggest that these transcription factors target different genes depending on the sex. Most of these sex-biased transcription factors were specific to a single tissue **(Figure 5B)**. However, six transcription factors (DLX2 (chr 2q31.1), ELK4 (chr 1q32.1), ETV3 (chr 1q23.1), KLF8 (chr Xp11.21), LHX6 (chr 9q33.2), and SOX14 (chr 3q22.3)) were identified in at least ten tissues, and 43 were identified in more than five tissues **(Figure S3)**.

**Figure 5.**
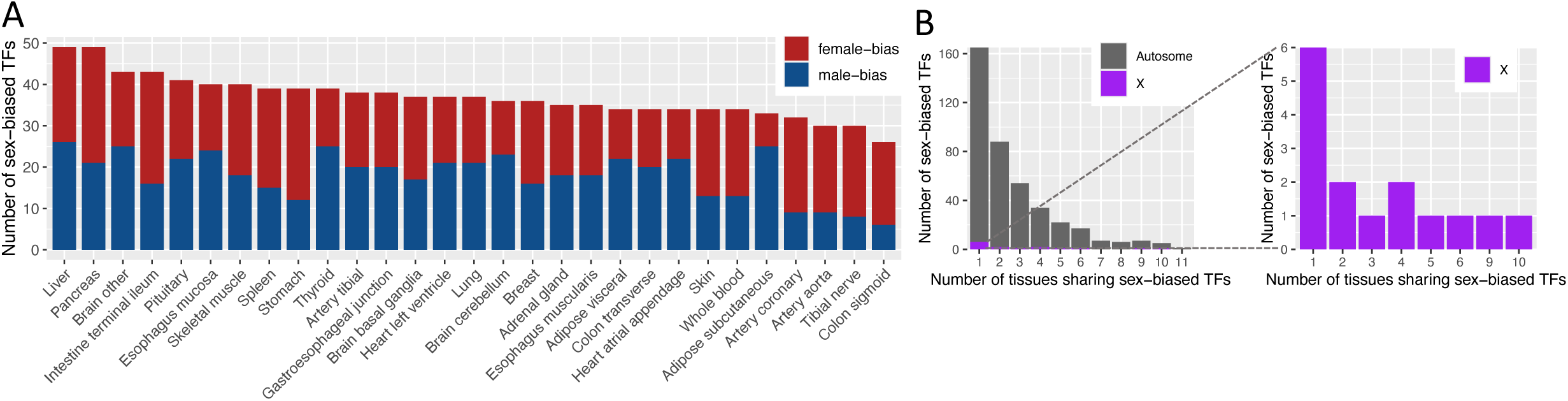
Sex-biased transcription factors are mostly tissue-specific. A) Number of sex-biased transcription factors (TFs) identified in each tissue. B) Distribution of the number of tissues that share sex-biased transcription factors, highlighting the number of autosomal and X-linked transcription factors.

We further examined motifs in the promoter region of these 43 transcription factors (3 on the X chromosome) identified as sex-biased in more than five tissues (**Figure 6, Table S4)**. Motifs for the sex hormones ESR1/ESR2 and AR were found to be significantly enriched in the promoter regions of these transcription factors, more than would be expected by chance (Fisher’s Exact test odds ratio 3.4 and 3.5; p-value 0.006 and 0.04 for ESR1 and AR, respectively). RARA, PPARA, TFAP2C, and KLF8 contain ESR1/ESR2 motifs and tend to target genes more strongly in females than in males across tissues. Transcription factors with AR motifs display varying patterns: TCF4 and HOXA13 target genes more strongly in females, while DLX5 and ARID2 target genes more strongly in males. While these sex-biased transcription factors did not show enrichment on the X chromosome beyond expected levels, the genes they differentially targeted were indeed enriched on the X chromosome (53% of the genes), particularly those that escape X chromosome inactivation (44% of the genes) (Fisher’s Exact test p<5×10^-39^).

**Figure 6.**
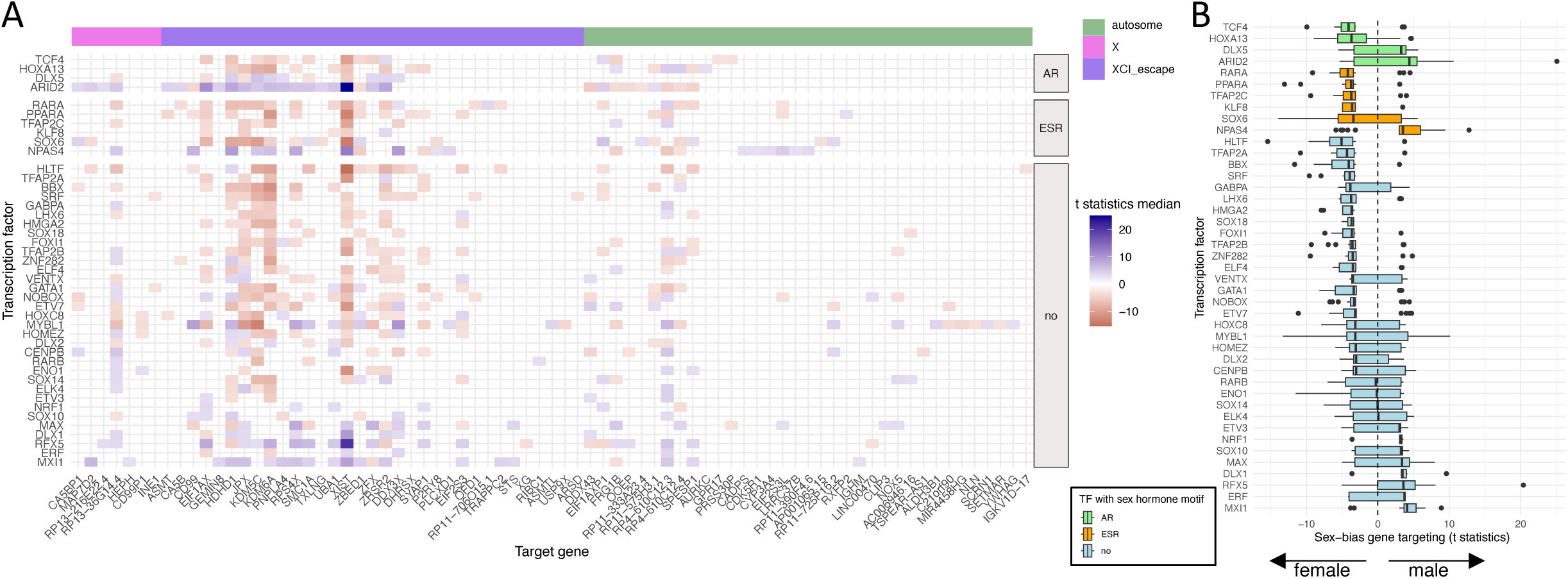
Sex-biased transcription factors across multiple tissues. A) Heatmap showing transcription factors identified as sex-biased in more than five tissues. For each gene, stronger sex-differential transcription factor targeting (absolute median t-statistic > 3 across tissues) is indicated in blue for males and red for females. The color of each heatmap cell reflects the median limma t-statistic for the sex difference in transcription factor–gene edge weights, calculated across tissues where the transcription factor was identified as sex-biased. B) Boxplots illustrating the distribution of t-statistics for genes differentially targeted by these sex-biased transcription factors (absolute median t-statistic > 3). Transcription factors with AR motifs in their promoters are shown in green, those with ESR1 or ESR2 motifs in orange, and all others in blue. Figure S3 displays the transcription factor targeting by tissue.

Thus, sex-biased transcription factors are largely tissue-specific and differentially target genes in males and females. The transcription factors that are consistently found to be sex-biased across multiple tissues are enriched for steroid hormone motifs and tend to target genes that escape X chromosome inactivation.

### Sex bias regulation of cancer genes in lung

We selected lung tissue as a representative example, as lung cancer remains the leading cause of cancer-related mortality worldwide and exhibits a higher incidence among younger females than males, as well as in females versus males without a prior smoking history^13,14^. Cancer genes were more strongly connected in the regulatory networks of female normal lung tissue compared to male, as measured by higher node in-degree and closeness **(Figure 7A)**. We identified sex-biased transcription factors, including those known to be cancer-associated such as JUNB and FOS. To identify the pathways enriched for genes targeted by these sex-biased transcription factors, we performed pre-ranked gene set enrichment analysis for each transcription factor by ranking their target genes based on sex-differential targeting. The pathways targeted by these transcription factors exhibited significant differences between males and females **(Figure 7B)**. Females showed a higher targeting of several cancer-related pathways, such as WNT, NOTCH, MAPK, and p53. In contrast, males exhibited a higher targeting of DNA repair pathways. These findings are consistent with previous lung cancer studies reporting higher mutation rates of *TP53* in females and a more efficient DNA repair mechanism in males^15–18^.

**Figure 7.**
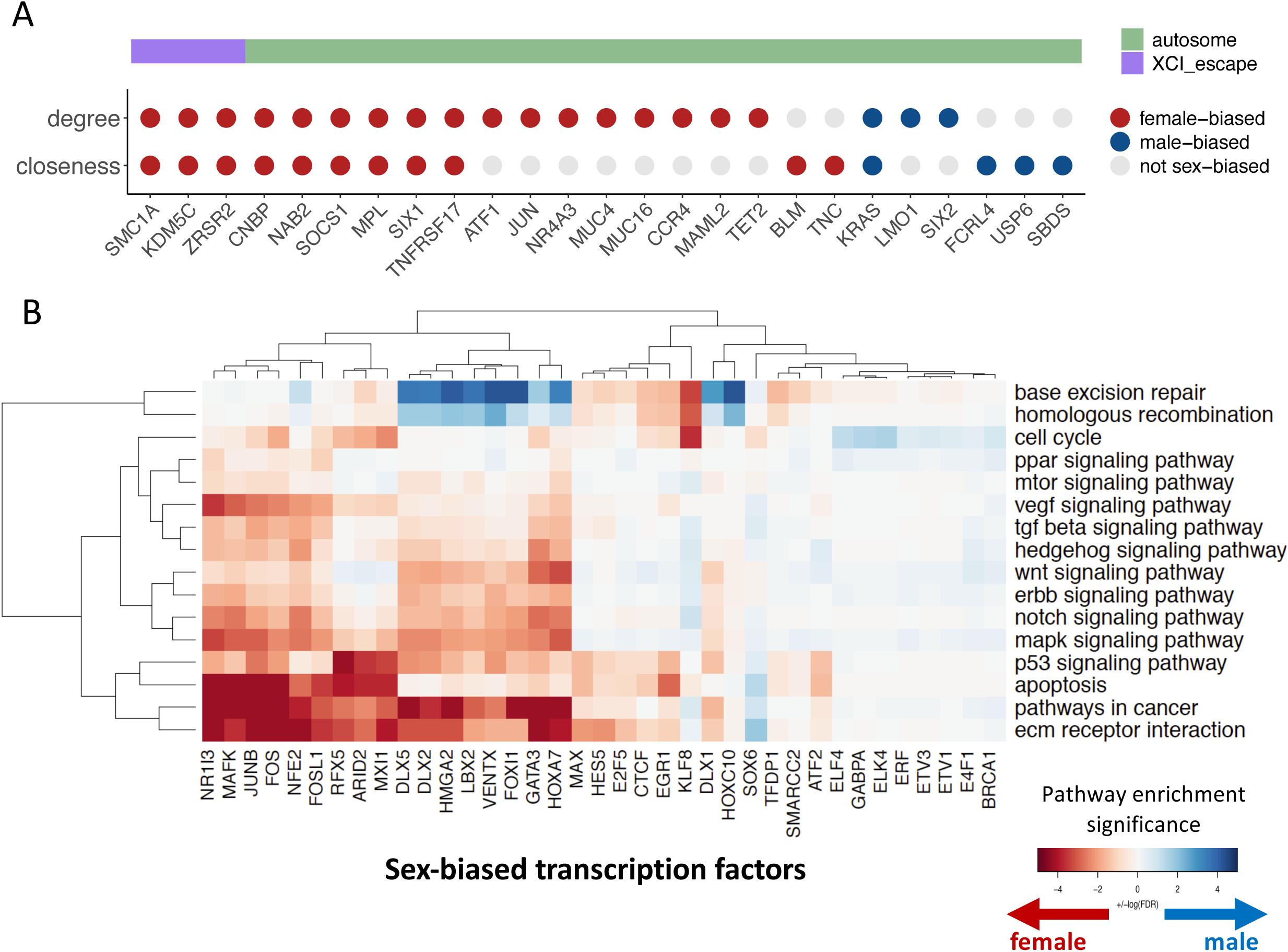
Transcription factors differentially target cancer-related genes and pathways in the lungs of males and females. A) In normal lung tissue from females, most cancer genes have higher transcription factor targeting (higher in-degree) and a shorter distance to all other genes in the network (higher closeness) compared to males. B) Cancer-related pathways targeted by sex-biased transcription factors in the lung. Colors indicate the significance of preranked gene set enrichment analysis performed for each transcription factor, based on the sex-differential targeting of their target genes.

We validated the sex-differential targeting of cancer-related pathways using an independent lung tissue dataset. We mined previously published gene regulatory networks from 315 lung tissue control samples in the Lung Tissue Research Consortium (LTRC) dataset^19^ and confirmed that in this independent dataset females have higher targeting of cancer-related pathways compared to males. These pathways included MAPK, p53, apoptosis, pathways in cancer, and extracellular matrix (ECM) receptor interaction (GSEA FDR<0.05). Validation in an independent dataset supports our approach and further indicates that cancer-related pathways are highly targeted in female lung tissue, which may help explain the higher risk of lung cancer observed in females compared to males.

## Discussion

By analyzing sex-differential gene regulatory networks across normal tissues, we have provided insights into sex-biased transcriptional mechanisms that may help explain observed differences in cancer incidence and etiology between males and females. Key signaling pathways, including WNT, NOTCH, and p53, exhibit differential transcriptional targeting across normal tissues based on sex. We found an overrepresentation of sex-biased cancer genes on the X chromosome, particularly among those escaping X chromosome inactivation. Our findings highlight the role of transcriptional regulation in sex differences in cancer incidence and emphasize the need to consider sex-specific gene regulatory processes when identifying targets for cancer prevention.

In previous research, we demonstrated that transcription factors exhibit sex-biased targeting of genes in colon^9^ and lung^10^ tumors, which is associated with differences in prognosis and therapeutic response. Here, we found that many transcription factors exhibit sex-biased targeting across normal tissues as well, which means that they target different genes involved in tumorigenesis in males and females even before cancer development. Most sex-biased transcription factors are tissue-specific. Sex-biased transcription factors that are consistently observed across multiple tissues are enriched for sex hormone response elements in their promoters, and they tend to target genes that escape X chromosome inactivation. This suggests that sex-biased transcription factors can influence sex differences in cancer susceptibility by differentially regulating cancer-related biological processes in a sex-specific manner.

The inherent dosage inequality of X chromosomes between males and females is typically balanced by the inactivation of one X chromosome in females. However, a subset of genes escapes this X chromosome inactivation, resulting in higher dosage in females compared to males^12^. Significantly, while 1.6% of genes across all autosomes are classified as cancer genes, this number rises to 6% for genes that escape X chromosome inactivation. Our study highlights an overrepresentation of sex-biased cancer genes that escape X chromosome inactivation, underscoring the importance of escape genes in cancer genetics, which is often overlooked as allosomes are not always included in genomics studies. We found that X-linked tumor suppressor genes such as *DDX3X*, *KDM5C*, *SMC1A*, and *ZRSR2*, which escape X chromosome inactivation, exhibit higher regulatory targeting in females. This elevated targeting suggests a potential protective mechanism against tumorigenesis, as females benefit from the expression of functional alleles from both X chromosomes. This observation aligns with the “Escape from X-Inactivation Tumor Suppressor” (EXITS) hypothesis, which suggests that females have a reduced risk for certain cancers due to the biallelic expression of EXITS genes, requiring a second genetic hit for inactivation. Supporting this hypothesis, Dunford et al. reported lower mutation rates of EXITS tumor tissues from females compared to males^20^. Despite these insights, the X chromosome has largely been excluded from Genome-Wide Association Studies (GWAS), limiting our understanding of its role in sex-biased cancer risk and incidence^21,22^. Future studies should prioritize the inclusion of the X chromosome in GWAS and other genomic investigations to gain insights into sex-specific genetic and genomic risk factors in cancer.

The sex-biased regulatory pattern of cancer-related pathways was evident across all tissues analyzed; however, the significance and direction of these patterns were highly tissue-specific. In tissues with a higher cancer incidence among females—such as breast, lung, and thyroid tissues—there was a marked increase in the targeting of cancer-related pathways in females. Conversely, in tissues with a higher cancer incidence in males—such as stomach, colon, and liver—there was a higher targeting in males. It has been well-established that dysregulation of key signaling pathways such as MAPK, NOTCH, WNT, and P53, all of which play crucial roles in cellular differentiation, proliferation, and survival, can lead to the development of cancer^23–26^. In this study, we have uncovered notable differences in the transcriptional regulation of these pathways between normal tissues of males and females. These sex-biased differences in gene regulation suggest that when mutations occur within these pathways, the resulting downstream effects may be influenced by the biological sex of the individual. Such differential regulation and mutation response could contribute to the observed sex differences in cancer susceptibility.

While our study provides valuable insights, it is important to note the limitations. We focused on 476 cancer genes as defined by the COSMIC Cancer Gene Census v92, and this may not represent all genes that drive cancer across every tissue type. We did not adjust the GTEx models for environmental exposures and lifestyle factors. We found associations between sex-biased transcription factor targeting and cancer incidence, but further research is required for causal conclusions. Additionally, because the GTEx dataset is composed primarily of White and African American individuals, larger studies involving more diverse populations are necessary to more extensively generalize our findings.

In conclusion, our analysis revealed significant sex-differential transcriptional targeting of cancer genes across normal tissues, with important implications for understanding sex differences in cancer incidence and identifying sex-specific targets for prevention. The increased targeting of tumor suppressor genes that escape X chromosome inactivation in females, along with the tissue-specific differences in signaling pathways associated with cancer development underscores the complex and tissue-dependent role of sex influencing cancer susceptibility. By shedding light on the role of the X chromosome and emphasizing the need for its inclusion in genomic studies, we pave the way for more comprehensive and sex-informed approaches to cancer research, including cancer, early diagnosis and treatment.

## Material and Methods

### Data

Sample-specific gene regulatory networks were previously inferred from RNA-sequencing data in the Genotype-Tissue Expression (GTEx) version 6.0 dataset^27^ using the PANDA^28^ and LIONESS^29^ algorithms, as described in Lopes-Ramos et al.^11^, and the networks are stored in the Gene Regulatory Network Database (GRAND)^30^. These networks capture transcription factor-gene regulatory relationships, where edge weights indicate the likelihood of a transcription factor regulating a target gene. To identify sex-biased regulatory interactions, a linear regression model was applied to detect edges with significantly different weights between males and females as reported in ^11^. Genes were classified into three categories: 1) male-biased genes (the proportion of male-biased edges is greater than 0.6); 2) female-biased genes (the proportion of male-biased edges is less than 0.4); and 3) sex-divergent genes (the proportion of male-biased edges is between 0.4 and 0.6), indicating similar numbers of male- and female-biased edges but distinct transcription factor regulators between sexes.

### Sex-differential network analysis

#### Sex-biased cancer gene targeting

We tested whether cancer genes were enriched for sex-biased regulatory edges relative to non-cancer genes. For this analysis, we defined cancer genes as 476 genes (237 oncogenes and 239 tumor suppressor genes) from the COSMIC Cancer Gene Census v92, of which 23 are located on the X chromosome (see **Table S2)**. These cancer genes were compared to a background of 29,767 non-cancer genes. For each chromosome, we quantified the proportion of cancer genes exhibiting sex-biased regulation. We then focused on the 1,018 genes on the X chromosome, classifying them according to four X chromosome inactivation (XCI) status categories as described by Tukiainen et al., 2017^12^: escape from inactivation (n=82), variable escape (n = 89), inactive (n = 388), and unknown (n = 459).

#### Pathways differentially targeted by sex

The gene targeting score, equivalent to the node in-degree, was calculated as the sum of all edge weights pointing to the gene. To compare the gene targeting scores between males and females for all autosomal and X chromosome genes (excluding Y chromosome genes), we used a linear regression model and the limma R package. The model was adjusted for potential confounders including race, age, body mass index (BMI), batch, and RNA Integrity Number (RIN). All genes (excluding Y chromosome genes) were ranked based on their t-statistics derived from the linear regression analysis and used as input for a pre-ranked gene set enrichment analysis (GSEA) using the fgsea R package, with KEGG pathways as the reference gene sets (downloaded from the Human MSigDB Collections KEGG v6.2). Pathways with an FDR<0.05 were considered differentially targeted by sex.

#### Network closeness and degree centrality

We filtered the networks to retain only the top 15% of edge weights. Closeness centrality was then calculated using the closeness function from the igraph R package, with mode = “all” to measure shortest paths while ignoring edge direction. A higher closeness centrality value indicates a node is connected to all other nodes in the network via shorter path lengths, suggesting greater reachability and potentially faster information flow throughout the network. We calculated each gene in-degree as the sum of regulatory edge weights connecting the gene with transcription factors.

#### Sex-biased transcription factors

To identify cascading regulation and capture potential indirect effects of transcription factors characteristic of master regulators, we applied the Independent Cascade Model^31,32^ to tissue- and sex-specific networks that included the top 15% of edge weights. Belief propagation (BP)^32,33^ was used to calculate, for each transcription factor, a score reflecting the number and weighted strength of its target genes within the network. To identify sex-biased transcription factors, we calculated the difference in rank scores between females and males, and classified transcription factors as sex-biased if their absolute z-score difference exceeded 2.

### Validation in an independent lung tissue dataset

To validate sex-biased targeting in the lung, we used sample-specific gene regulatory networks that we had previously generated from the Lung Tissue Research Consortium (LTRC) dataset, dbGap accession phs001662^19^. For the current study, we focused on networks derived from individuals with normal lung function as determined by spirometry. We had applied the same differential targeting analysis to compare gene targeting scores between males and females, followed by a pre-ranked gene set enrichment analysis using KEGG pathways^34^. Pathways with an FDR<0.05 were considered differentially targeted by sex.

### Data sharing statement

Gene regulatory networks using the GTEx data are available through Gene Regulatory Network Database (GRAND)^30^; https://grand.networkmedicine.org/tissues. The Lung Tissue Research Consortium study data are available from the National Center for Biotechnology Information Database of Genotypes and Phenotypes (dbGaP) under accession no. phs001662.

## Acknowledgments

Figure 1 was created in BioRender; Lopes Ramos, C. (2025) https://BioRender.com/sayx3vm

## Author Contributions

CML-R: Conceptualization, Data curation, Data analysis, Interpretation, Writing, Funding acquisition; RB: Conceptualization, Data analysis, Interpretation, Writing; MBG: Conceptualization, Interpretation, Writing; VF: Conceptualization, Interpretation, Writing; ES: Conceptualization, Interpretation, Writing; KHS: Conceptualization, Interpretation, Writing; KG: Conceptualization, Interpretation, Writing, Supervision; JQ: Conceptualization, Interpretation, Writing, Funding acquisition, Supervision; DLD: Conceptualization, Interpretation, Writing, Funding acquisition, Supervision.

## Supplemental Figures

**Supplemental Figure 1.**
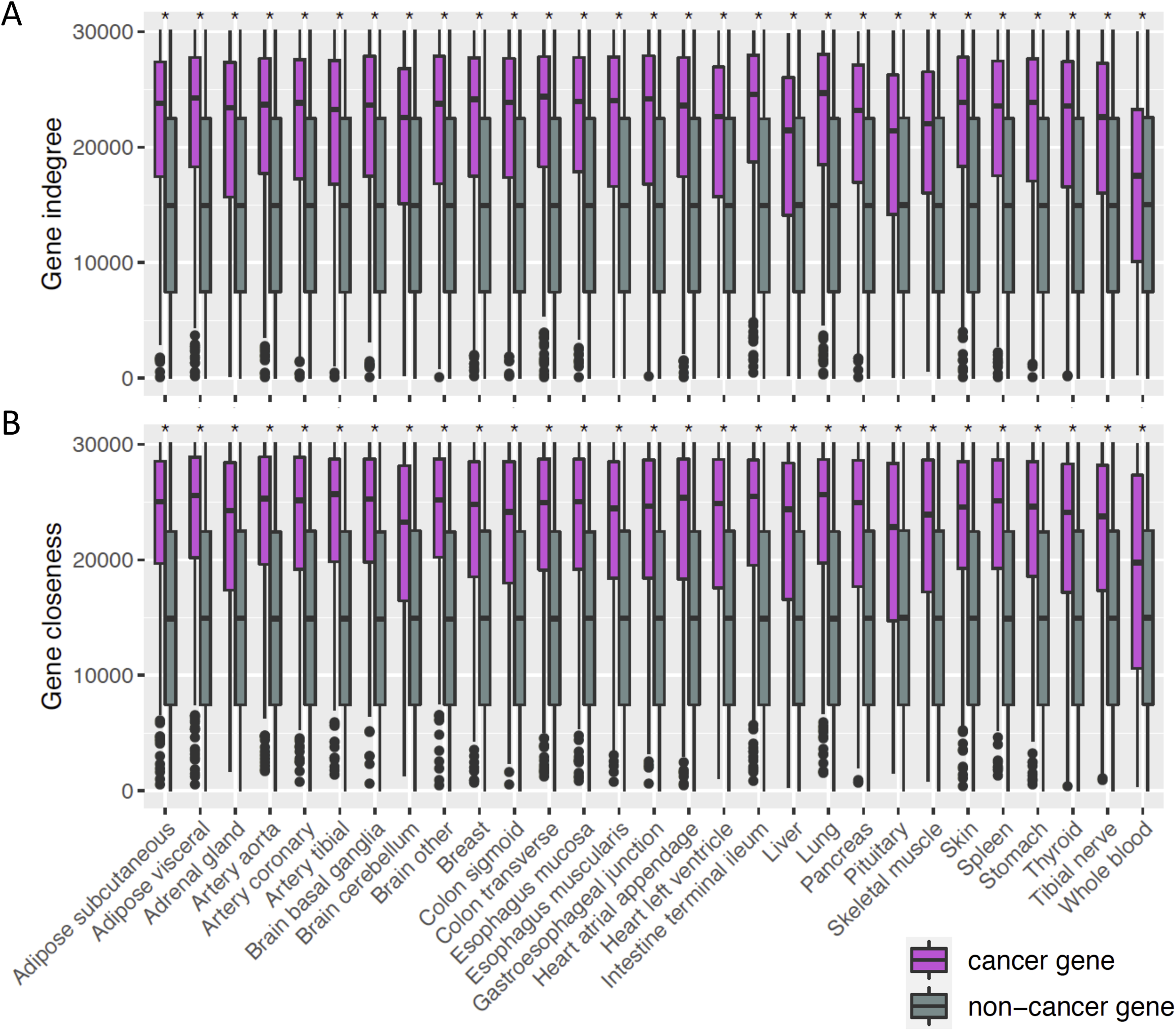
Cancer genes have higher regulatory influence in the gene regulatory networks than non-cancer genes. Plots show rank distribution of the network centralities in-degree and closeness. A) Cancer genes have higher transcription factor targeting (higher in-degree) compared to non-cancer genes. B) Cancer genes have shortest distance to all other genes in the networks (higher closeness) compared to non-cancer genes. * p<0.05, t-test

**Supplemental Figure 2.**
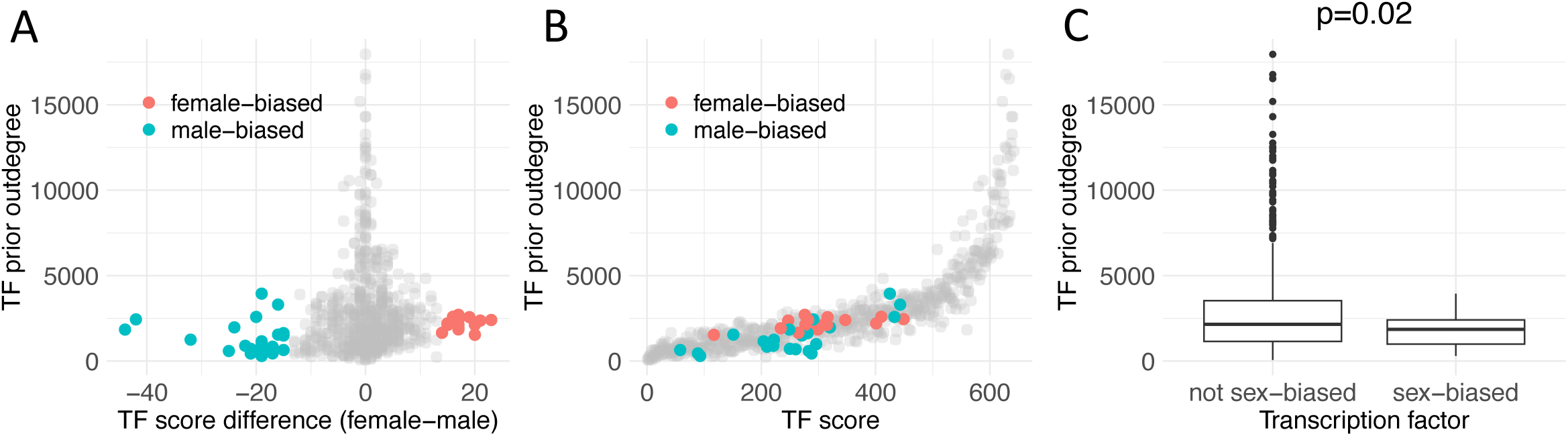
Using cascade model to identify sex-biased transcription factors in lung. **A)** Scatterplot of transcription factor prior out-degree (the number of genes containing the transcription factor’s motif in the network prior) versus the ranked score difference between sexes. For each sex- and tissue-specific gene regulatory network, the cascade model calculates a ranked score representing the likelihood that a transcription factor can regulate the majority of target genes. The difference in ranked scores between males and females is shown on the x-axis. Female-biased transcription factors (ranked score difference z-score > 2) are highlighted in red, and male-biased transcription factors (z-score < −2) are shown in blue. **B)** Scatterplot displaying the relationship between transcription factor prior out-degree and ranked scores for transcription factors in both male and female lung networks. Transcription factors with significant sex bias (as in panel A) are highlighted. **C)** Boxplot comparing the distribution of prior out-degree between identified sex-biased transcription factors and transcription factors without sex bias (p-value from a Mann-Whitney test). These plots represent values specifically for lung tissue. Similar results were observed across other tissues.

**Supplemental Figure 3.**
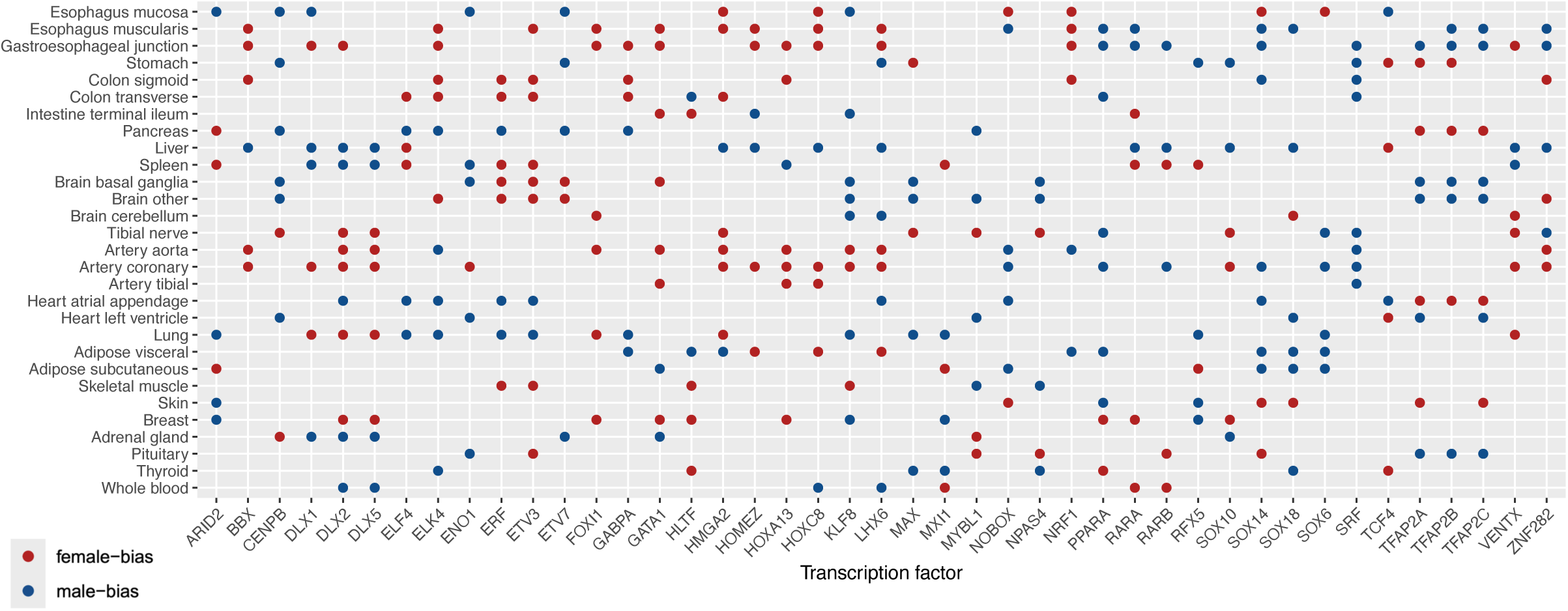
Sex-biased transcription factors from the cascade model identified in more than five tissues.

## Supplemental Table

**Table S1**. Sample characteristics by sex. P-values compare male and female groups within each tissue (Chi-square test for categorical variables; t-test for continuous variables).

**Table S2.** List of the cancer gene set used in this study, which includes 237 oncogenes and 239 tumor suppressor genes, totaling 476 genes that are expressed in the GTEx dataset. Gene types (oncogene or tumor suppressor) were defined using the COSMIC Cancer Gene Census v92.

**Table S3.** KEGG pathways enriched for genes differentially targeted by sex. Pre-ranked gene set enrichment analysis in each tissue was performed on genes ranked by sex differential targeting. Positive normalized enrichment scores (NES) indicate higher targeting in females and negative NES indicate higher targeting in males.

**Table S4.** Sex-biased transcription factors identified in more than five tissues.

| TF | Number of tissues | Chromosome |
| --- | --- | --- |
| DLX2 | 11 | 2 |
| ELK4 | 10 | 1 |
| ETV3 | 10 | 1 |
| KLF8 | 10 | X |
| LHX6 | 10 | 9 |
| SOX14 | 10 | 3 |
| DLX5 | 9 | 7 |
| ERF | 9 | 19 |
| GATA1 | 9 | X |
| HMGA2 | 9 | 12 |
| PPARA | 9 | 22 |
| TFAP2A | 9 | 6 |
| TFAP2C | 9 | 20 |
| CENPB | 8 | 20 |
| HOXC8 | 8 | 12 |
| SOX18 | 8 | 20 |
| SRF | 8 | 6 |
| TFAP2B | 8 | 6 |
| ZNF282 | 8 | 7 |
| ARID2 | 7 | 12 |
| DLX1 | 7 | 2 |
| HOXA13 | 7 | 7 |
| MYBL1 | 7 | 8 |
| NOBOX | 7 | 7 |
| RARA | 7 | 17 |
| VENTX | 7 | 10 |
| BBX | 6 | 3 |
| ELF4 | 6 | X |
| ENO1 | 6 | 1 |
| ETV7 | 6 | 6 |
| FOXI1 | 6 | 5 |
| GABPA | 6 | 21 |
| HLTF | 6 | 3 |
| HOMEZ | 6 | 14 |
| MAX | 6 | 14 |
| MXI1 | 6 | 10 |
| NPAS4 | 6 | 11 |
| NRF1 | 6 | 7 |
| RARB | 6 | 3 |
| RFX5 | 6 | 1 |
| SOX10 | 6 | 22 |
| SOX6 | 6 | 11 |
| TCF4 | 6 | 18 |

## References

1. Tosakoon, S., Lawrence, W. R., Shiels, M. S. & Jackson, S. S. Sex Differences in Cancer Incidence Rates by Race and Ethnicity: Results from the Surveillance, Epidemiology, and End Results (SEER) Registry (2000–2019). Cancers (Basel) 16, (2024).

2. Jackson, S. S., Marks, M. A., Katki, H. A., Cook, M. B., Hyun, N., Freedman, N. D., Kahle, L. L., Castle, P. E., Graubard, B. I. & Chaturvedi, A. K. Sex disparities in the incidence of 21 cancer types: Quantification of the contribution of risk factors. Cancer 128, 3531–3540 (2022).

3. Dong, M., Cioffi, G., Wang, J., Waite, K. A., Ostrom, Q. T., Kruchko, C., Lathia, J. D., Rubin, J. B., Berens, M. E., Connor, J. & Barnholtz-Sloan, J. S. Sex Differences in Cancer Incidence and Survival: A Pan-Cancer Analysis. Cancer Epidemiol Biomarkers Prev 29, 1389–1397 (2020).

4. Selvaraj, R. C., Cioffi, G., Waite, K. A., Jackson, S. S. & Barnholtz-Sloan, J. S. A Pan-Cancer Analysis of Age and Sex Differences in Cancer Incidence and Survival in the United States, 2001–2020. Cancers (Basel) 17, (2025).

5. Rubin, J. B. The spectrum of sex differences in cancer. Trends Cancer 8, 303–315 (2022).

6. Ozga, M., Nicolet, D., Mrózek, K., Yilmaz, A. S., Kohlschmidt, J., Larkin, K. T., Blachly, J. S., Oakes, C. C., Buss, J., Walker, C. J., Orwick, S., Jurinovic, V., Rothenberg-Thurley, M., Dufour, A., Schneider, S., Sauerland, M. C., Görlich, D., Krug, U., Berdel, W. E., Woermann, B. J., Hiddemann, W., Braess, J., Subklewe, M., Spiekermann, K., Carroll, A. J., Blum, W. G., Powell, B. L., Kolitz, J. E., Moore, J. O., Mayer, R. J., Larson, R. A., Uy, G. L., Stock, W., Metzeler, K. H., Grimes, H. L., Byrd, J. C., Salomonis, N., Herold, T., Mims, A. S. & Eisfeld, A. K. Sex-associated differences in frequencies and prognostic impact of recurrent genetic alterations in adult acute myeloid leukemia (Alliance, AMLCG). Leukemia 38, 45 (2023).

7. Li, C. H., Haider, S., Shiah, Y.-J., Thai, K. & Boutros, P. C. Sex Differences in Cancer Driver Genes and Biomarkers. Cancer Res 78, 5527–5537 (2018).

8. Yuan, Y., Liu, L., Chen, H., Wang, Y., Xu, Y., Mao, H., Li, J., Mills, G. B., Shu, Y., Li, L. & Liang, H. Comprehensive Characterization of Molecular Differences in Cancer between Male and Female Patients. Cancer Cell 29, 711–722 (2016).

9. Lopes-Ramos, C. M., Kuijjer, M. L., Ogino, S., Fuchs, C. S., DeMeo, D. L., Glass, K. & Quackenbush, J. Gene Regulatory Network Analysis Identifies Sex-Linked Differences in Colon Cancer Drug Metabolism. Cancer Res 78, 5538–5547 (2018).

10. Saha, E., Ben Guebila, M., Fanfani, V., Fischer, J., Shutta, K. H., Mandros, P., DeMeo, D. L., Quackenbush, J. & Lopes-Ramos, C. M. Gene regulatory networks reveal sex difference in lung adenocarcinoma. Biol Sex Differ 15, (2024).

11. Lopes-Ramos, C. M., Chen, C.-Y., Kuijjer, M. L., Paulson, J. N., Sonawane, A. R., Fagny, M., Platig, J., Glass, K., Quackenbush, J. & DeMeo, D. L. Sex Differences in Gene Expression and Regulatory Networks across 29 Human Tissues. Cell Rep 31, 107795 (2020).

12. Tukiainen, T., Villani, A.-C., Yen, A., Rivas, M. A., Marshall, J. L., Satija, R., Aguirre, M., Gauthier, L., Fleharty, M., Kirby, A., Cummings, B. B., Castel, S. E., Karczewski, K. J., Aguet, F., Byrnes, A., Lappalainen, T., Regev, A., Ardlie, K. G., Hacohen, N. & MacArthur, D. G. Landscape of X chromosome inactivation across human tissues. Nature 550, 244–248 (2017).

13. Ragavan, M. & Patel, M. I. The evolving landscape of sex-based differences in lung cancer: a distinct disease in women. European Respiratory Review 31, (2022).

14. Siegel, R. L., Kratzer, T., Giaquinto, A. N., Sung, H. & Jemal, A. Cancer statistics, 2025. CA Cancer J Clin 75, 10 (2025).

15. Kure, E. H., Ryberg, D., Hewer, A., Phillips, D. H., Skaug, V., Bæera, R. & Haugen, A. *p53* mutations in lung tumours: relationship to gender and lung DNA adduct levels. Carcinogenesis 17, 2201–2205 (1996).

16. Guinee, D. G., Travis, W. D., Trivers, G. E., De Benedetti, V. M., Cawley, H., Welsh, J. A., Bennett, W. P., Jett, J., Colby, T. V & Tazelaar, H. Gender comparisons in human lung cancer: analysis of p53 mutations, anti-p53 serum antibodies and C-erbB-2 expression. Carcinogenesis 16, 993–1002 (1995).

17. Ryberg, D., Hewer, A., Phillips, D. H. & Haugen, A. Different Susceptibility to Smoking-induced DNA Damage among Male and Female Lung Cancer Patients. Cancer Res 54, 5801–5803 (1994).

18. Qingyi Wei, Q., Cheng, L., Amos, C. I., Wang, L.-E., Guo, Z., Hong, W. K. & Spitz, M. R. Repair of Tobacco Carcinogen-Induced DNA Adducts and Lung Cancer Risk: a Molecular Epidemiologic Study. J Natl Cancer Inst 92, 1764–1772 (2000).

19. Lopes-Ramos, C. M., Shutta, K. H., Ryu, M. H., Huang, Y., Saha, E., Ziniti, J., Chase, R., Hobbs, B. D., Yun, J. H., Castaldi, P., Hersh, C. P., Glass, K., Silverman, E. K., Quackenbush, J. & DeMeo, D. L. Sex-biased Regulation of Extracellular Matrix Genes in COPD. Am J Respir Cell Mol Biol 72, (2024).

20. Dunford, A., Weinstock, D. M., Savova, V., Schumacher, S. E., Cleary, J. P., Yoda, A., Sullivan, T. J., Hess, J. M., Gimelbrant, A. A., Beroukhim, R., Lawrence, M. S., Getz, G. & Lane, A. A. Tumor-suppressor genes that escape from X-inactivation contribute to cancer sex bias. Nat Genet 49, 10–16 (2017).

21. Sun, L., Wang, Z., Lu, T., Manolio, T. A. & Paterson, A. D. eXclusionarY: 10 years later, where are the sex chromosomes in GWASs? Am J Hum Genet 110, 903–912 (2023).

22. Khramtsova, E. A., Wilson, M. A., Martin, J., Winham, S. J., He, K. Y., Davis, L. K. & Stranger, B. E. Quality control and analytic best practices for testing genetic models of sex differences in large populations. Cell 186, 2044–2061 (2023).

23. Braicu, C., Buse, M., Busuioc, C., Drula, R., Gulei, D., Raduly, L., Rusu, A., Irimie, A., Atanasov, A. G., Slaby, O., Ionescu, C. & Berindan-Neagoe, I. A Comprehensive Review on MAPK: A Promising Therapeutic Target in Cancer. Cancers (Basel) 11, 1618 (2019).

24. Shi, Q., Xue, C., Zeng, Y., Yuan, X., Chu, Q., Jiang, S., Wang, J., Zhang, Y., Zhu, D. & Li, L. Notch signaling pathway in cancer: from mechanistic insights to targeted therapies. Signal Transduction and Targeted Therapy 2024 9:1 9, 1–37 (2024).

25. Zhan, T., Rindtorff, N. & Boutros, M. Wnt signaling in cancer. Oncogene 36, 1461–1473 (2017).

26. Wang, H., Guo, M., Wei, H. & Chen, Y. Targeting p53 pathways: mechanisms, structures and advances in therapy. Signal Transduction and Targeted Therapy 2023 8:1 8, 1–35 (2023).

27. Carithers, L. J., Ardlie, K., Barcus, M., Branton, P. A., Britton, A., Buia, S. A., Compton, C. C., Deluca, D. S., Peter-Demchok, J., Gelfand, E. T., Guan, P., Korzeniewski, G. E., Lockhart, N. C., Rabiner, C. A., Rao, A. K., Robinson, K. L., Roche, N. V., Sawyer, S. J., Segrè, A. V., Shive, C. E., Smith, A. M., Sobin, L. H., Undale, A. H., Valentino, K. M., Vaught, J., Young, T. R., Moore, H. M., Barker, L., Basile, M., Battle, A., Boyer, J., Bradbury, D., Bridge, J. P., Brown, A., Burges, R., Choi, C., Colantuoni, D., Cox, N., Dermitzakis, E. T., Derr, L. K., Dinsmore, M. J., Erickson, K., Fleming, J., Flutre, T., Foster, B. A., Gamazon, E. R., Getz, G., Gillard, B. M., Guigó, R., Hambright, K. W., Hariharan, P., Hasz, R., Im, H. K., Jewell, S., Karasik, E., Kellis, M., Kheradpour, P., Koester, S., Koller, D., Konkashbaev, A., Lappalainen, T., Little, R., Liu, J., Lo, E., Lonsdale, J. T., Lu, C., MacArthur, D. G., Magazine, H., Maller, J. B., Marcus, Y., Mash, D. C., McCarthy, M. I., McLean, J., Mestichelli, B., Miklos, M., Monlong, J., Mosavel, M., Moser, M. T., Mostafavi, S., Nicolae, D. L., Pritchard, J., Qi, L., Ramsey, K., Rivas, M. A., Robles, B. E., Rohrer, D. C., Salvatore, M., Sammeth, M., Seleski, J., Shad, S., Siminoff, L. A., Stephens, M., Struewing, J., Sullivan, T., Sullivan, S., Syron, J., Tabor, D., Taherian, M., Tejada, J., Temple, G. F., Thomas, J. A., Thomson, A. W., Tidwell, D., Traino, H. M., Tu, Z., Valley, D. R., Volpi, S., Walters, G. D., Ward, L. D., Wen, X., Winckler, W., Wu, S., Zhu, J., Abdallah, A., Addington, A., Anderson, J. M., Bender, P. K., Cosentino, M., Diaz-Mayoral, N., Engel, T., Garci, F., Green, A., Hammond, T., Jaffe, K., Keen, J., Kennedy, M., Kigonya, P., Lander, B., Nampally, S., Ny, C., Robb, J., Santhanum, V., Sharopova, N., Singh, S., Soria, C., Sturcke, A., Sukari, S., Thomson, E. J., Tomaszewski, M., Trowbridge, C., Udoye, F., Vanscoy, D., Vatanian, N., Wilder, E. L. & Williams, P. A Novel Approach to High-Quality Postmortem Tissue Procurement: The GTEx Project. Biopreserv Biobank 13, 311–317 (2015).

28. Glass, K., Huttenhower, C., Quackenbush, J. & Yuan, G.-C. Passing messages between biological networks to refine predicted interactions. PLoS One 8, e64832 (2013).

29. Kuijjer, M. L., Tung, M. G., Yuan, G., Quackenbush, J. & Glass, K. Estimating Sample-Specific Regulatory Networks. iScience 14, 226–240 (2019).

30. Ben Guebila, M., Lopes-Ramos, C. M., Weighill, D., Sonawane, A. R., Burkholz, R., Shamsaei, B., Platig, J., Glass, K., Kuijjer, M. L. & Quackenbush, J. GRAND: a database of gene regulatory network models across human conditions. Nucleic Acids Res 50, D610–D621 (2022).

31. Nalluri, J. J., Rana, P., Barh, D., Azevedo, V., Dinh, T. N., Vladimirov, V. & Ghosh, P. Determining causal miRNAs and their signaling cascade in diseases using an influence diffusion model. Sci Rep 7, 1–14 (2017).

32. Burkholz, R. & Quackenbush, J. Cascade Size Distributions: Why They Matter and How to Compute Them Efficiently. Proc AAAI Conf Artif Intell 8, 6840 (2021).

33. Burkholz, R. Efficient message passing for cascade size distributions. Sci Rep 9, 1–10 (2019).

34. Kanehisa, M., Sato, Y., Kawashima, M., Furumichi, M. & Tanabe, M. KEGG as a reference resource for gene and protein annotation. Nucleic Acids Res 44, D457–62 (2015).

